# Extremophilic Laccase-Mediator System Enabling Near-Complete Degradation of Doxycycline

**DOI:** 10.64898/2026.10.13.758911

**Authors:** Matías Donoso, Diego Muñoz-Gacitúa, Gabriel Salfate, Carla Toledo-Neira, Jenny M. Blamey

## Abstract

Using an extremophilic laccase with a phenolic redox mediator, we rapidly abolished doxycycline antibacterial activity and achieved near-complete chemical degradation (>97%) within 2 h under mild conditions. Antibiotic residues in aquatic and terrestrial environments drive the emergence and spread of microbial resistance genes, yet many treatment strategies are limited by slow kinetics or incomplete transformation. Here, we evaluated doxycycline degradation by the recombinant FNT laccase across pH and temperature conditions, comparing direct enzymatic oxidation with a laccase–mediator system. RP-HPLC quantified doxycycline removal. Residual toxicity was assessed using growth inhibition assays with *Bacillus subtilis* and *Escherichia coli* BL21. In the absence of a mediator, oxidation produced only partial loss of function, indicating that structural modification alone does not guarantee pharmacological inactivation. In contrast, the addition of acetosyringone markedly increased biodegradation efficiency, yielding >97% doxycycline depletion and eliminating antibacterial activity. To rationalize condition-dependent performance, we complemented experiments with molecular simulations of doxycycline binding at the laccase active site, revealing pH-dependent differences in stability and substrate interactions consistent with enhanced turnover under alkaline conditions. These results demonstrate that an extremophilic laccase–mediator system provides rapid, effective detoxification of doxycycline, supporting its potential for treatment of antibiotic-contaminated effluents.

## 1. Introduction

The development of antibiotics revolutionized modern medicine, turning once-lethal infections into manageable conditions. For instance, Staphylococcus aureus-induced pneumonia–once a death sentence with a 80% mortality rate in the pre-antibiotic era(Hofer, 2022)–now has a mortality rate of just ∼20%, largely related to rare, high-risk patient cohorts(Thabet et al., 2022). Antibiotics have thus not only extended life expectancy but have also transformed the management of infectious diseases worldwide.

However, the injudicious use and overconsumption of antibiotics have led to a concerning trend: *the same lifesaving drugs are accumulating in ecosystems at alarming rates*. Global antibiotic consumption has surged by 65% since 2000, with forecasts predicting a 200% rise by 2030(Hofer, 2022). Over 70% of consumed antibiotics are excreted unmetabolized by humans and livestock(Antos et al., 2024; De Ilurdoz et al., 2022). As a result, these compounds frequently enter municipal wastewater streams and are present in animal manure, which is often applied as fertilizer. Consequently, elevated concentrations of antibiotics are now regularly detected in surface waters, groundwater, and agricultural soils worldwide(Gros et al., 2021; Liu et al., 2022; Zeng et al., 2022).

The consequences of this antibiotic contamination pose significant ecological and health risks. The presence of antibiotics in ecosystems can drive the emergence of antibiotic resistance genes (ARGs)(Larsson & Flach, 2022), compromising the effectiveness of current and future treatments. Furthermore, antibiotics can disrupt nutrient cycling and decomposition processes, exemplified by studies showing denitrification inhibition in wastewaters and shallow lakes(Zhu et al., 2023). Moreover, exposure to antibiotics has been linked to toxicity in non-target organisms(Da Silva Freitas et al., 2022), potentially disrupting delicate ecological balances and impacting ecosystem services.

Among the antibiotics contributing to environmental contamination, tetracyclines represent a major class of broad-spectrum agents, effective against both gram-positive and gram-negative microorganisms. Due to their versatility, tetracyclines are frequently detected in municipal wastewater effluents(Miao et al., 2004), agricultural manure(Hamscher et al., 2002; Hu et al., 2010), and surface water bodies(Kolpin et al., 2002; Lindsey et al., 2001). Doxycycline, a second-generation tetracycline derivative, is particularly prevalent in these matrices. Its cost-effective synthesis, high clinical efficacy, and versatile applications (spanning human medicine, aquaculture, and intensive livestock farming, especially in developing regions) have driven widespread use. Consequently, residual doxycycline persists through waste streams, and the need arises to remove it from these environments.

Various strategies have been explored to mitigate antibiotic contamination in ecosystems, including chemical coagulation/flocculation, adsorption via activated charcoal or other adsorbents, physicochemical oxidation/degradation, and the use of biofilters, among others^3^. While these approaches have shown promise, they are not without drawbacks. They either are costly or do not neutralize a significant concentration of antibiotics. But perhaps most critically, these methods often fail to address the issue of robustness, as they are typically optimized for controlled laboratory settings rather than the harsh, dynamic environments in which they are ultimately deployed. This limitation underscores the need for more resilient and adaptable solutions, capable of withstanding the rigors of real-world applications.

The quest for more resilient and adaptable solutions has led researchers to explore alternative approaches, leveraging the unique properties of extremophilic organisms. These microorganisms have evolved to function optimally in extreme habitats–such as geothermal springs, hypersaline lakes, acidic mine drainages, and alkaline soda lakes–conferring them natural stability under conditions that would be lethal to mesophilic organisms. Enzymes derived from extremophiles have been shown to exhibit enhanced stability, activity, and selectivity, making them attractive candidates for industrial applications(Krüger et al., 2018; Niehaus et al., 1999; Rawat et al., 2024). For instance, thermophilic enzymes have been used in biofuel production(Jiang et al., 2017), while psychrophilic enzymes have found applications in food processing and bioremediation(Parvizpour et al., 2021). Similarly, alkalophilic enzymes, which thrive in high-pH environments, have been explored for their potential in wastewater treatment and bioremediation(Borkar, 2015; Johnson et al., 2024). The discovery of these novel enzymes has opened up new avenues for the development of sustainable and efficient industrial processes.

Among extremozyme candidates for bioremediation, laccases (multicopper oxidases; EC 1.10.3.2) warrant particular attention due to their versatile substrate promiscuity. These enzymes catalyse the oxidation of diverse phenolic compounds with preferential efficiency toward diphenolic structures(Espina et al., 2021). This versatility is further enhanced when laccases are used in conjunction with redox mediators, forming laccase-mediator systems (LMS), which enable the indirect oxidation of non-phenolic substrates through radical chain reactions. This mechanism enables LMS to degrade recalcitrant environmental pollutants, including pharmaceutical antibiotics, efficiently. One such laccase of interest is the **FNT laccase**–isolated from a thermoalkaliphilic *Bacillus sp*.(Atalah et al., 2018) –which exemplifies this potential with exceptional biocatalytic resilience. It maintains structural integrity and activity across extensive thermal (20-90 °C) and pH (6.0-9.0) ranges, inheriting stability from its extremophilic origin. Notably, FNT laccase demonstrated efficacy in tetracycline-class antibiotic degradation when paired with acetosyringone (AS)(Sánchez-SanMartín et al., 2024), a naturally abundant phenolic mediator derived from renewable lignin sources. This combination offers an economically viable and environmentally compatible degradation platform.

Building upon its demonstrated resilience and tetracycline degradation efficiency, the FNT laccase system represents a promising biocatalytic platform for mitigating antibiotic contamination in environmentally relevant conditions. To comprehensively evaluate its bioremediation potential for doxycycline, this study deployed an LMS, composed by the FNT laccase and AS, to achieve complete biodegradation of aqueous doxycycline solutions. Degradation efficiency was quantified via high-performance liquid chromatography (HPLC), while functional attenuation of antibiotic activity was verified through growth inhibition assays against model Gram-positive (*Bacillus subtilis*) and Gram-negative (*Escherichia coli*) cultures. Molecular simulations were performed to rationalize the observed effects. The results of this study have significant implications for the development of sustainable and effective bioremediation strategies, highlighting the potential of FNT laccase as a valuable tool for mitigating the environmental impact of antibiotic pollutants.

## 2. Methodology

### 2.1 Enzyme assay

A stock solution of 20 mg/mL of the antibiotic doxycycline (DOX) was prepared, which was then dissolved in 1800 µL of HPLC-grade water and 200 µL of 0.1 M HCl(Council of Europe, 2023). In addition, a stock solution of 10 mM redox mediator AS was prepared and resuspended in HPLC-grade methanol. For the enzyme, a stock solution with a concentration of 10 mg/mL was prepared from lyophilized laccase resuspended in HPLC-grade water.

The reactions were performed in triplicate in a volume of 1000 µL and consisted of 0.1 mg/mL of enzyme and 0.5 mg/mL of antibiotic in presence or absence of 0.5 mM AS redox mediator. Each reaction was brought to a final volume by adding a 20 mM phosphate buffer, which was adjusted with sodium hydroxide to the required pH (6, 7, and 8) and diluted in HPLC-grade water at temperatures of 25, 32, and 40°C. Each reaction was incubated for 2 hours at 180 rpm at the corresponding temperature. Once the time had elapsed, the reactions were stopped by adding an equal volume of acetonitrile at −20°C and vortexing for 30 seconds.

### 2.2 Quantification of the biodegradation of doxycycline

To evaluate the oxidation of the antibiotic obtained under the procedure described in the previous section, reverse-phase high-performance liquid chromatography (RP-HPLC) was used. Before chromatographic analysis, the samples obtained from the experiments were filtered with a 0.22 µm PTFE filter and stored at 4°C until analysis. At each tested condition, two control experiments were performed, in one the antibiotic was incubated in the absence of laccase and mediator, and in another the antibiotic was incubated in the absence of laccase but in the presence of the mediator. Degradation of the antibiotic was not observed in any case where the laccase was not added to the mix.

Reverse-phase liquid chromatography was performed on a Shimadzu LC-20AT HPLC system with a C-18 Phenomenex reverse-phase column (particle size 2.6 µm, length 150 mm, internal diameter 4.6 mm) under isocratic conditions of 25% ACN, 70% TFA 0.1%, and 5% methanol at 40°C.

Chromatography was performed at 1 mL/min, with an injection volume of 20 µL. The chromatograms of the samples were obtained using an SPD-M20A diode array detector. The antibiotic was measured at a wavelength of 250 nm. The areas were analyzed using the equipment’s integrated software. The percentage of antibiotic degradation was calculated as follows:

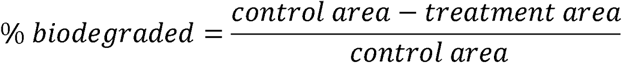

### 2.3 Microorganism culturing

The microorganisms *Escherichia coli* BL21 (E. coli BL21) was cultured in LB medium (10 g/L tryptone, 5 g/L yeast extract, 10 g/L NaCl) at pH 7 and 30°C, and *Bacillus subtilis* (B. subtilis) was cultivated in HVMOD-D1 medium (48 g/L NaCl, 5 g/L glucose, 4 g/L yeast extract, 4 g/L peptone, 3.5 g/L casamino acids, 2 g/L sodium citrate, 4 g/L MgSO_4_ x 7H_2_O, 0.05 g/L CaCl_2_, 0.4 g/L K_2_HPO_4_, 0.04 g/L Fe(NH_4_)_2_(SO_4_)_2_ x 6H_2_O) at pH 6.5 and 30°C. All cultures were grown in an aerobic medium at 180 rpm.

The microorganism *Escherichia coli BL21* (*E. coli* BL21) was cultured in LB medium (10 g/L tryptone, 5 g/L yeast extract, 10 g/L sodium chloride) at pH 7 and 30 °C, while *Bacillus subtilis* (*B. subtilis*) was cultivated in HVMOD-D1 medium (48 g/L sodium chloride, 5 g/L glucose, 4 g/L yeast extract, 4 g/L peptone, 3.5 g/L casamino acids, 2 g/L sodium citrate, 4 g/L magnesium sulphate heptahydrate, 0.05 g/L calcium chloride, 0.4 g/L dipotassium hydrogen phosphate, 0.04 g/L ammonium iron(II) sulphate hexahydrate) at pH 6.5 and 30 °C. All cultures were grown under aerobic conditions at 180 rpm.

### 2.4 Analysis of residual toxicity after biodegradation

To determine residual toxicity, diffusion plate tests were performed with LB agar medium pH 7 and HVMOD-D1 agar pH 6.5 using the microorganisms mentioned above. Each microorganism was cultured to an OD of 0.1 at 600 nm, using the previously sterilized liquid medium as a target. Each plate was inoculated with 150 µL of the culture, and a lawn was grown on the solid medium. Then, following the modified Kirby-Bauer disc diffusion method (Biemer, 1973; Boyle et al., 1973), 6 mm diameter filter paper discs previously sterilized for 30 minutes with UV light were added to the inoculated agar. 5 µL of the reaction mixtures with and without redox mediator, incubated for 2 hours, were added to the sterilized discs. As controls for the assay, the antibiotics were incubated in the absence of laccase under the same conditions described above. The plates were incubated at 30°C for 18 hours to observe the diameter of the inhibition halo. The inhibition zone halos were measured using the ImageJ program (Wayne Rasband, NIH).

### 2.5 Molecular simulation analysis

3D structures of the FNT laccase enzyme were obtained via homology modeling(Waterhouse et al., 2018). Structures of the antibiotics were generated from their SMILES sequences, which were obtained from the PubChem database, using OpenBabel(O’Boyle et al., 2011). Molecular docking using Autodock Vina(Eberhardt et al., 2021) was performed to obtain a structure of the ligand docked to the active site of the enzyme. P2rank(Krivák & Hoksza, 2018) was used to programmatically find and characterize the active site of the enzyme. pH was simulated through adjusting the protonation states of each amino acid in the enzyme; this was calculated using Propka(Olsson et al., 2011), which performs pKa predictions on each residue based on the chemical environment around each one.

Doxycycline parameters were obtained using the OpenFF Sage forcefield specification, which is compatible with AMBER biopolymer forcefields(Boothroyd et al., 2023). Their atomic partial charges were calculated from a Merk-Kollman scheme, which was performed on a single point calculation using the Hartree-Fock method; a basis set of 6-311(d,p) was picked for this last calculation, which was performed using Gaussian 16(Frisch et al., 2016). Hamiltonian Monte-Carlo (HMC) simulations were performed using the docking results as the initial structure. Each evolution of the Markov chain consisted of a displacement of the ligand in a random direction, up to 2.5 Å, followed by a random rotation of the ligand and completing the iteration with a short molecular dynamics simulation of 100 ps. The chain was evolved 250 times, or until the interaction energies could be fitted to a Boltzmann distribution function (p<0.05). The molecular dynamics calculations were performed with a timestep of 2 fs, and a Langevin integrator under a BAOAB splitting(Kieninger & Keller, 2022). Each stochastic update of velocities (the *O* step in the splitting) simulated an interaction with a heat bath at 305 K. Water was simulated using an OBC2 implicit solvent model. Enzyme was simulated using the AMBER14 force field(Maier et al., 2015).

Interaction energy (Δ*E*_inter_) between the antibiotic and the enzyme was calculated from the following thermodynamic cycle:

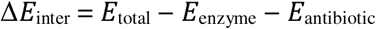

Where *E*_total_ is the energy of the full system, *E*_enzyme_ is the energy of the enzyme without the antibiotic present, and *E*_antibiotic_ is the opposite.

### 2.6 Statistical analysis

All tests were performed in triplicate and included their respective controls. Each result was considered to be significant only if there was a statistically significant difference from its control, as per a student’s t-test (p<0.05).

## 3. Results

### 3.1 Effect of pH and temperature on doxycycline activity reduction

The effect of pH and temperature on the enzymatic reduction of doxycycline antibacterial activity was first evaluated.

Except for the tests at 40 °C, all controls, which were incubations without the enzyme, resulted in no degradation of the antibiotic. At 40 °C, a significant fraction of the doxycycline was decomposed after two hours of incubation without the addition of FNT laccase. We decided not to continue with the enzymatic assays at this temperature, as the enzyme would not have made a difference. The success of the oxidation mediated by the enzyme was associated with a change of colour in the solution; they acquired a reddish-brown colour, and no precipitation was observed after two hours of incubation.

At temperatures of 25°C and 32°C, after two hours of treating doxycycline with FNT laccase, a reduction in antibiotic activity is observed; the magnitude of this effect depends on the pH at which the incubation was performed. About 16±10% of the antibiotic activity was inhibited at pH 6 and 7. There was not a statistically significant difference between these pH values (p-value: 0.08). However, there was a significant increase at pH 8 (p-value: 0.008), where about 26% of the antibiotic activity was inhibited. Incubation temperature did not seem to make a significant difference, and the remaining antimicrobial activity seemed to be equally effective against *B. subtilis* as against *E. coli*.

**Figure 1:**
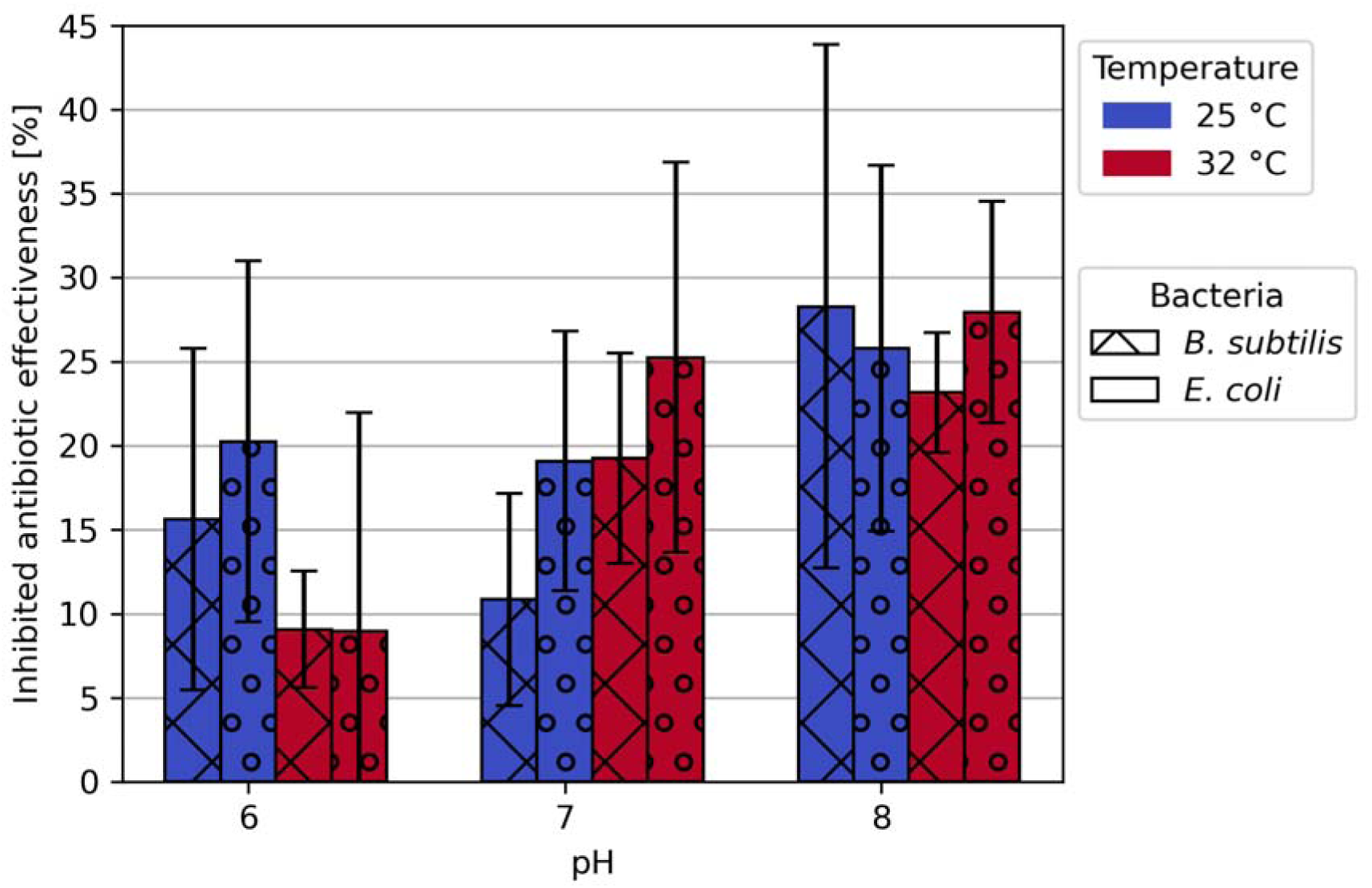
Inhibited antibiotic effectiveness of doxycycline after treatment with FNT laccase using *B. subtilis* (crossed bars) and *E. coli* (dotted bars). Experiments performed at 25 °C and 32 °C.

### 3.2 Doxycycline biodegradation quantified by HPLC

To further quantify doxycycline transformation, biodegradation was assessed by RP-HPLC. When assessing the fraction of doxycycline that was biodegraded by the laccase enzyme, the results showed that at pH 6 and 7 similar results were obtained via growth inhibition tests: there is about a 6±3% of the doxycycline being biodegraded with no statistically significant differences between both conditions. However, at pH 8, about 80±2% of the doxycycline was measured to be degraded. At first glance, this result would not match what was obtained via growth inhibition tests, where only ∼16% of the antibiotic activity was inhibited.

The reasoning behind this discrepancy will be discussed below. A temperature increment from 25 C to 32 °C results in slightly more degradation, about 3% more in all tested pH values. This apparent discrepancy suggests that structural modification alone does not necessarily account for the loss of biological activity.

**Figure 2:**
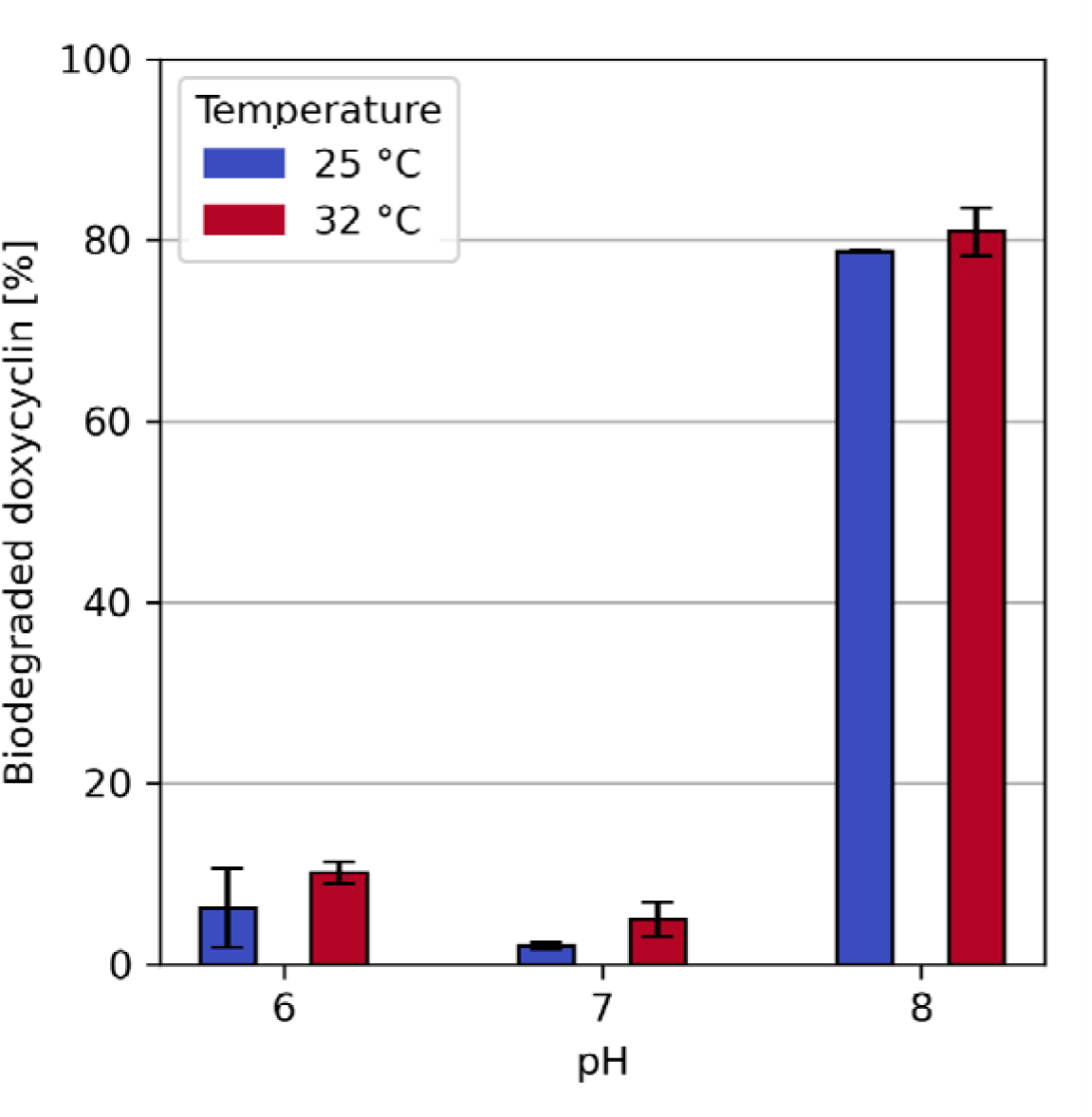
Fraction of doxycycline biodegraded after treatment with FNT laccase. Comparison between different pH and temperature used during treatment.

### 3.3 Effect of the redox mediator (AS) on degradation efficiency

The addition of AS significantly improves the fraction of the doxycycline that is biodegraded by the enzyme at both temperatures tested, reaching 97.1±0.3% and 98.5±0.7% of the antibiotic degraded at 25 °C and 32 °C, respectively. These results confirm the critical role of the mediator in achieving near-complete degradation.

**Figure 3:**
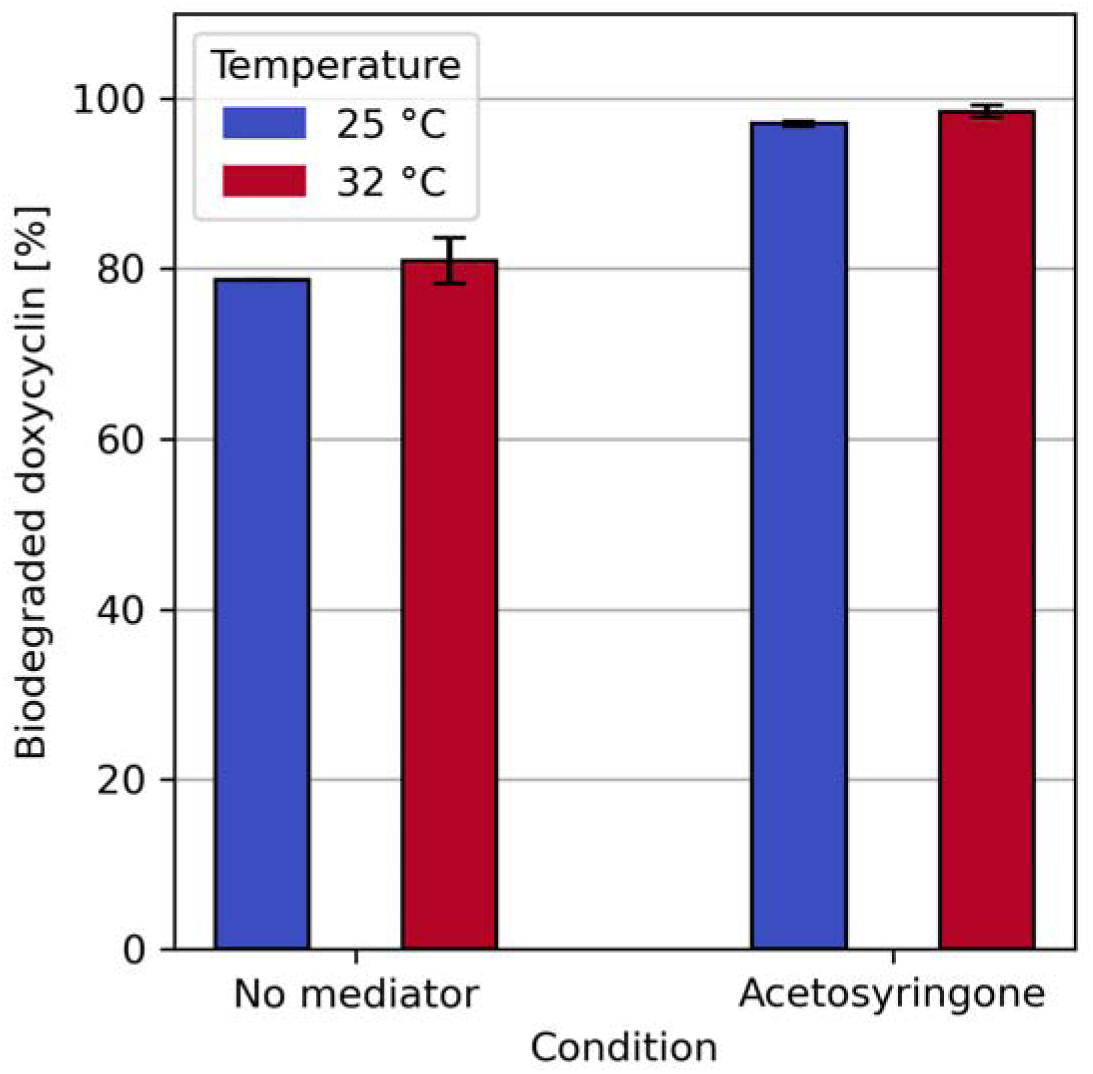
Fraction of the doxycycline that is biodegraded after treatment with FNT laccase. Comparison between treatment without a redox mediator and the addition of AS. Experiments performed at pH 8, temperatures of 25 °C and 32 °C.

### 3.4 Elimination of antibacterial activity in the laccase–mediator system

This improvement is further observed when the results of the growth inhibition tests are performed. Upon the addition of AS, doxycycline loses virtually all its toxicity towards *B. subtilis* and *E. coli*, at both temperatures tested.

**Figure 4:**
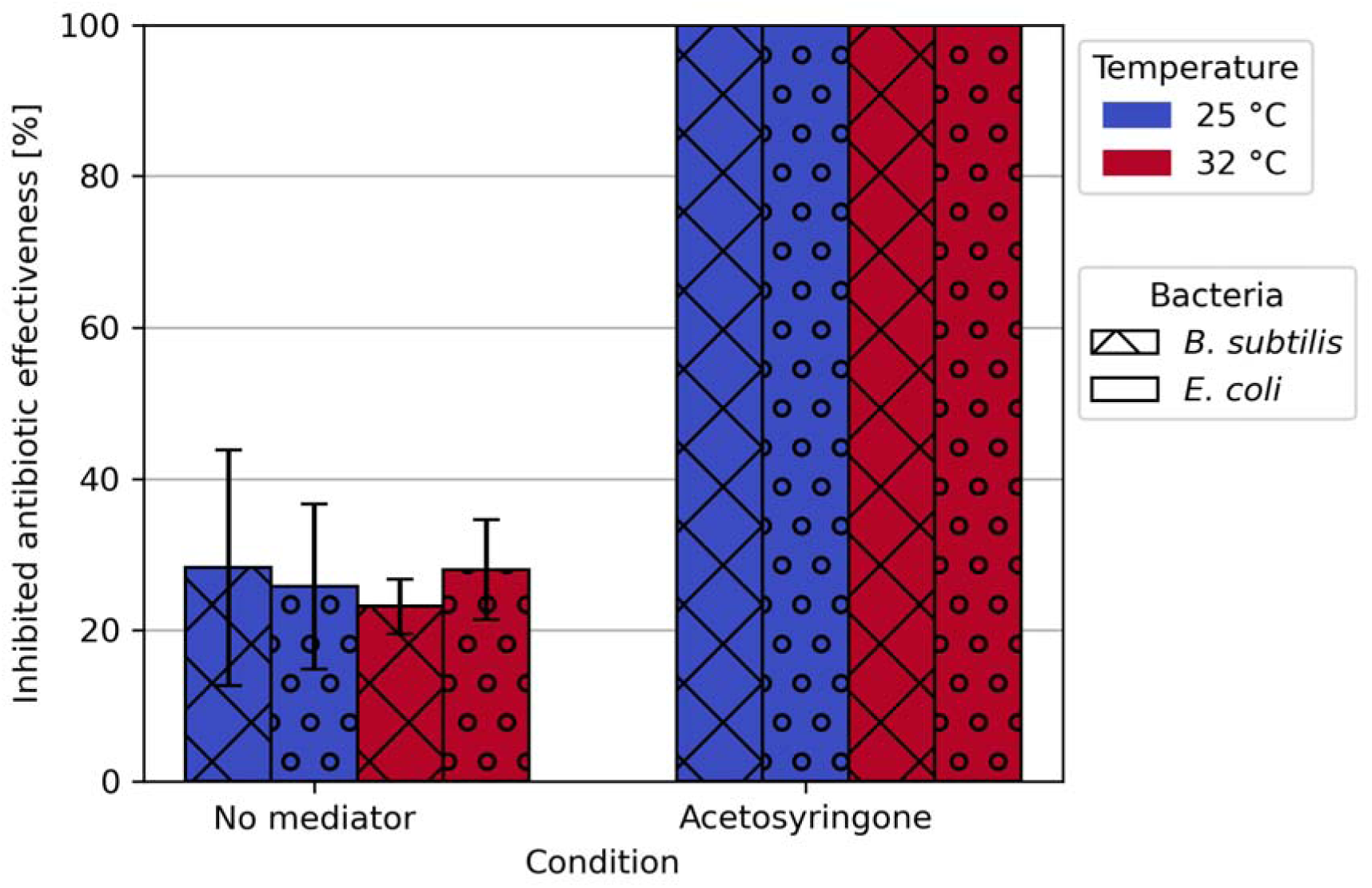
Inhibited antibiotic effectiveness of doxycycline after treatment with FNT laccase, comparison between treatment with no redox mediator and AS addition. Using *B. subtilis* (crossed bars) and *E. coli* (dotted bars). Experiments performed at 25 °C and 32° C.

### 3.5 Molecular simulation analysis of enzyme–substrate interactions

Molecular simulations of a doxycycline molecule interacting with the active site of laccase enzyme, show that, at the equilibrium, there is not a significant correlation between the distance of doxycycline from the active site at equilibrium and its experimental activity. The non-oxidized doxycycline stabilizes 7.3 Å away from the active site, which is closer than at pH 6, where it stabilizes at 11.5 Å from the active site. This would suggest that doxycycline has a more favourable interaction with the laccase enzyme at pH 7 than at pH 6, but this is not what is reflected in the experiment, where we observe a lesser effectivity at pH 7. This situation changes when one adds simulations of the oxidized doxycycline (that is, with a missing electron) into consideration, where we observe that, at pH values of 6.0 and 8.0, doxycycline stabilizes further away from the active site, whereas at pH 7, the antibiotic stays in place after being oxidized. This is clearer in Figure 6, where at pH 7, there is a negligible change in interaction after the substrate has been oxidized.

**Figure 5:**
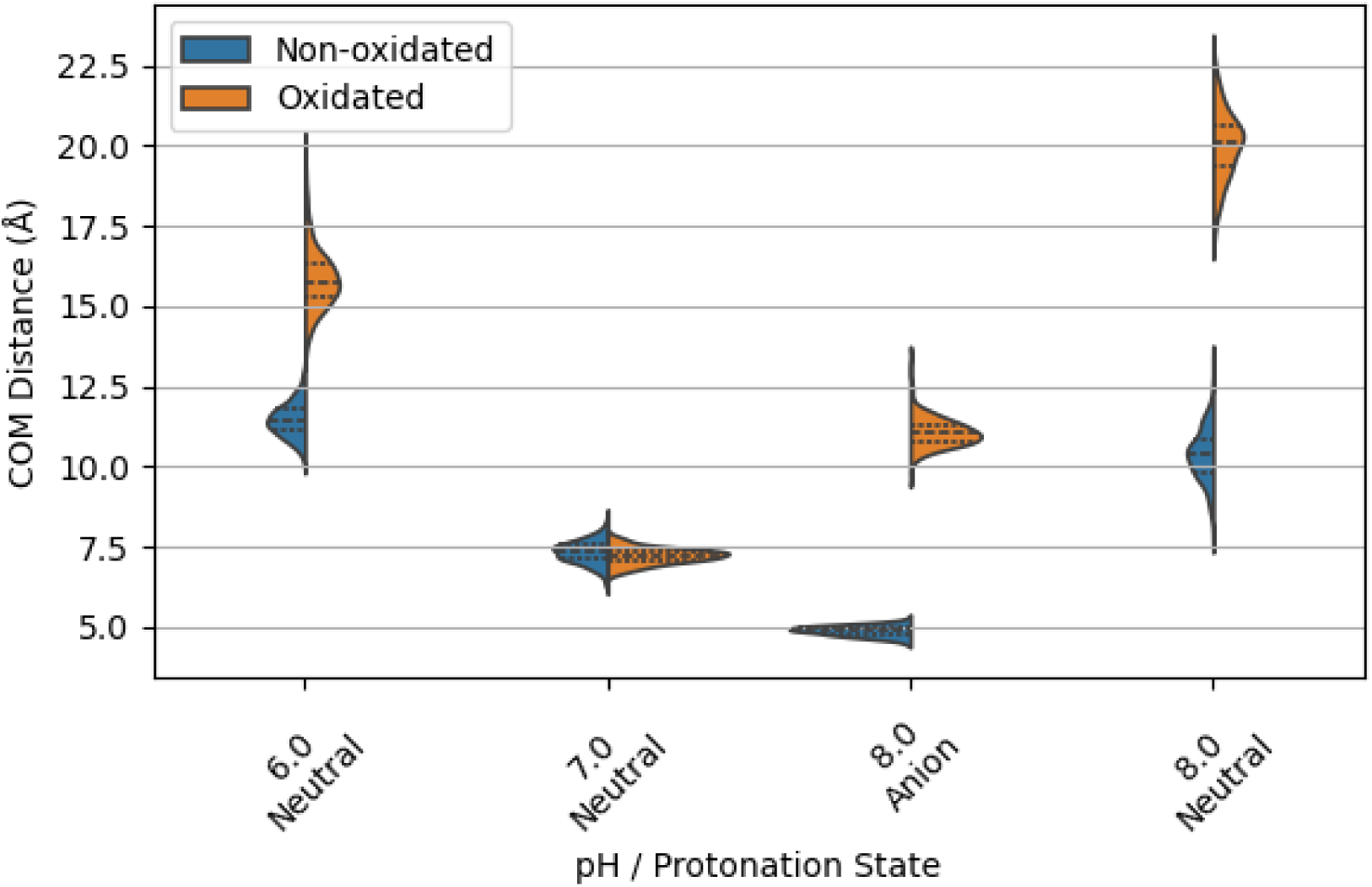
Calculated distance between the doxycycline center-of-mass (COM) and T1 copper atom of the active site of laccase enzyme. Comparison between non-oxidized and oxidized states of the antibiotic.

**Figure 6:**
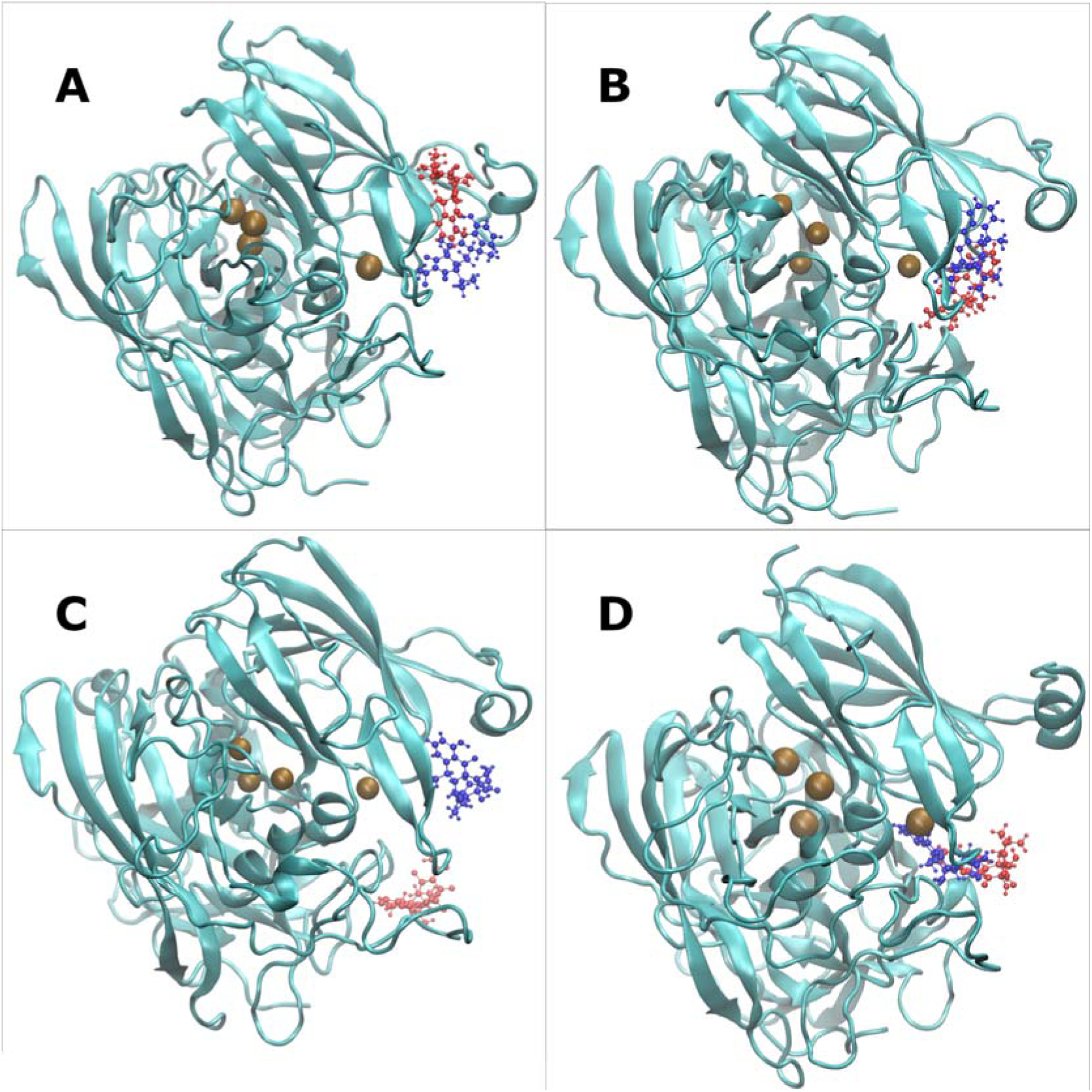
Doxycycline molecule interacting with FNT laccase at different pH and doxycycline protonation states. FNT laccase is shown as a ribbon diagram with its copper atoms shown in brown colour. Doxycycline is shown in colour blue when not oxidized and in red when oxidized. (A) Laccase at pH 6, zwitterionic doxycycline. (B) Laccase at pH 7, zwitterionic doxycycline. (C) Laccase at pH 8, zwitterionic doxycycline. (D) Laccase at pH 8, anionic doxycycline.

**Figure 7:**
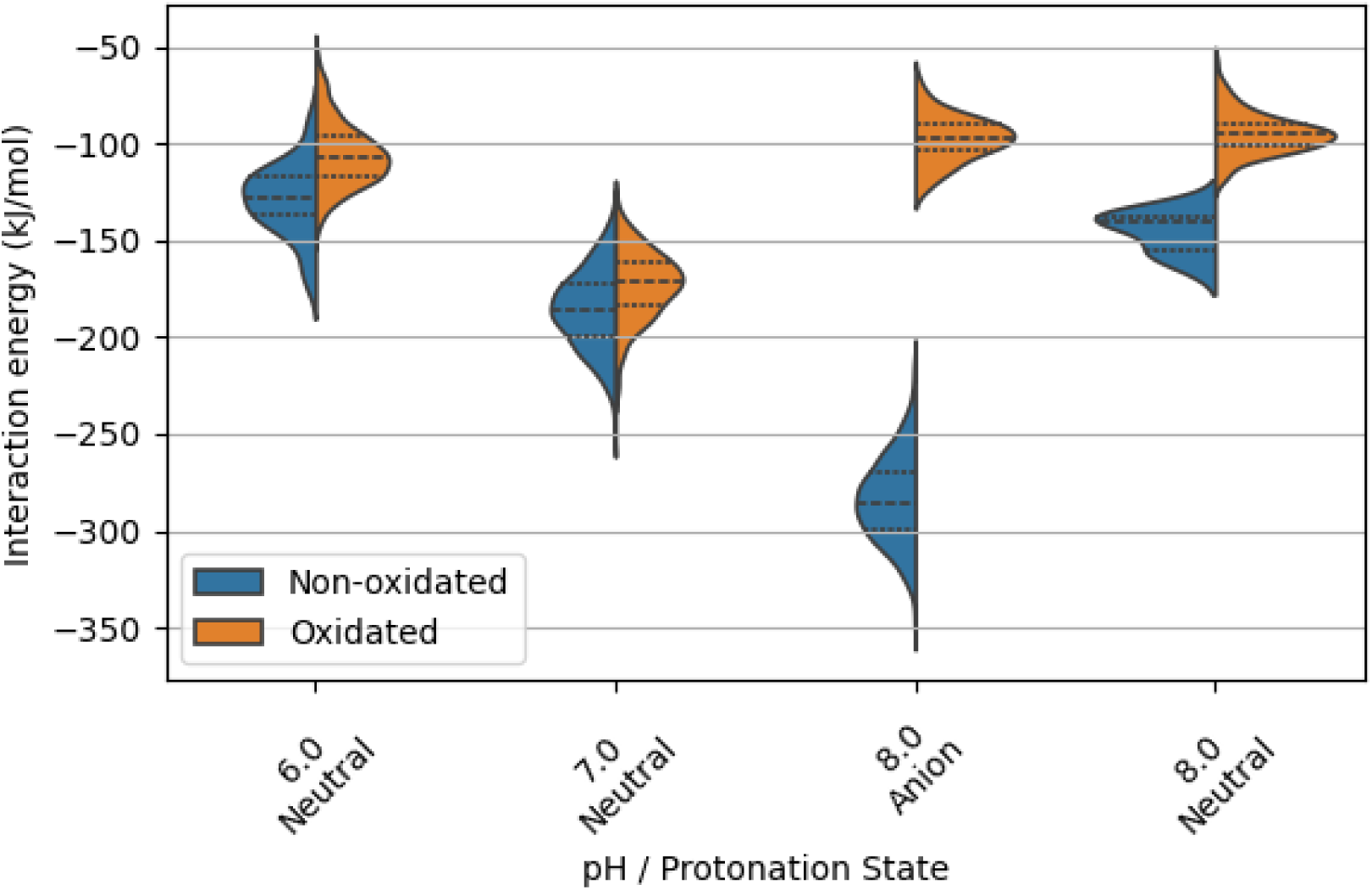
Calculated interaction energy between doxycycline and FNT laccase enzyme at the active site. Results from HMC simulations.

**Figure 8:**
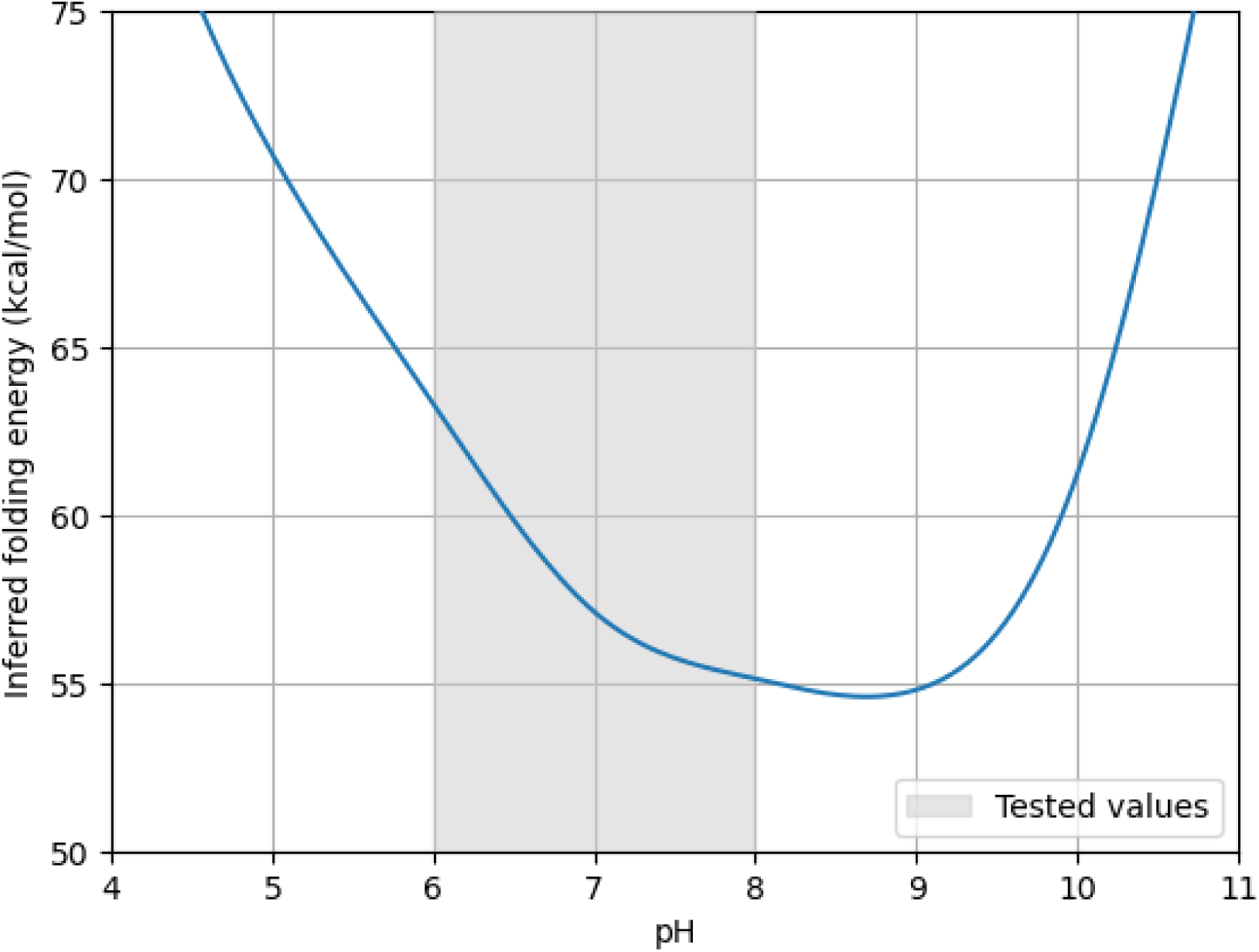
ML-inferred folding energy of FNT laccase enzyme at different pH values. The range that was experimentally tested is shaded in grey.

Regarding the interaction energy of the doxycycline and the enzyme, at pH 6 and 7, the oxidized doxycycline has a similar interaction strength with the active site as its non-oxidized counterpart. This is not the case at pH 8, where the oxidized doxycycline, which not only stabilises further away from the active site, but also has a lower interaction energy than its non-oxidized form.

## 4. Discussion

Our results demonstrate a critical distinction between partial structural modification and complete functional degradation of doxycycline by FNT laccase. In the absence of redox mediator AS, enzymatic oxidation yielded oxidized derivatives, visually confirmed by solution darkening that predominantly retained antibiotic activity against both Gram-positive *(B. subtilis)* and Gram-negative (*E. coli*) bacteria. Crucially, the limited bioactivity reduction (16-26%) despite significant molecular oxidation implies that conventional laccase catalysis frequently fails to disrupt doxycycline’s core pharmacophore. This pharmacophore resilience would explain the persistent antibiotic potency observed where most primary oxidation products likely retain this structural motif.

This limitation is overcome by the AS-mediated laccase system. This addition expanded degradation efficacy (>97%) and completely inhibited antibiotic function. We postulate that AS radicals, generated by FNT laccase, act as *cascades* that attack non-phenolic moieties or unstable sites *outside the core pharmacophore*. This fundamentally alters the degradation mechanism: instead of targeted phenolic oxidation, the LMS enables broad *scission of the critical molecular architecture* responsible for antimicrobial action. Consequently, residue fragmentation progresses beyond compromised pharmacophores to bio-inert fragments.

Regarding the observed pH-dependent efficacy, the causes can be attributed to multiple factors, but two key aspects stand out. First, inferred folding energy calculations (see figure 8) reveal that the enzyme’s stability is enhanced at higher pH values, with a maximum stability at pH 8.7. This is consistent with the alkaliphilic nature of the organism from which the enzyme was isolated.

Secondly, the molecular simulations indicate that the oxidized doxycycline products stabilise further away from the active site at pH 8, compared to lower pH values. This thermodynamic preference for clearance of the oxidation product allows for a more efficient catalytic cycle, as the enzyme can more readily bind to another doxycycline molecule. Furthermore, interaction energy calculations reveal that, at pH 6 and 7, the oxidized doxycycline has a similar interaction energy with the active site compared to its non-oxidized form, whereas at pH 8, this interaction becomes weaker after oxidation. This weakened interaction likely facilitates the clearance of the active site, enabling the enzyme to process more substrate molecules efficiently. This observation is especially true at pH 8, where anionic doxycycline, which should be about two-thirds of the species at that pH, not only has the strongest interaction with FNT laccase, but also has the biggest difference in interaction when compared with its oxidized form (ΔE_interaction_ = +186 kJ/mol). We believe this is a major contributing factor in making this condition have a higher activity.

The combined effects of the increased enzyme stability, optimized product clearance, and reduced interaction energy at pH 8 likely contribute to the observed increased efficacy of

## 5. Conclusions

Our study demonstrates that the combination of FNT laccase and AS offers a solution for the efficient biodegradation and structural annihilation of doxycycline. The approach not only rapidly degrades the antibiotic but also effectively destroys its pharmacophore.

Notably, our results show that FNT laccase achieves this degradation in just two hours at ambient temperature, outperforming previously reported methods that require at least 24 hours at high temperature to achieve the same result(Han et al., 2023). This highlights the potential of laccase-mediator systems as a rapid and efficient solution for antibiotic degradation.

The importance of destroying the pharmacophore of doxycycline cannot be overstated, as it guarantees complete loss of biological activity rather than partial transformation into less active derivatives. Otherwise, the risks associated with antibiotic contamination persist.

Our findings suggest that laccase-mediator systems have significant potential for wastewater treatment, pharmaceutical effluent processing, and soil remediation. By leveraging the enzymatic activity of laccase and the mediator AS, these systems offer a promising approach for degrading antibiotics and other contaminants.

Our molecular simulation studies provide valuable insights into the mechanisms underlying the laccase-mediated degradation of doxycycline. From our calculations, we propose that effective doxycycline oxidation by a laccase enzyme requires three key conditions: first, the oxidized substrate should thermodynamically stabilize away from the active site to allow another molecule to be oxidized; second, the interaction energy of the oxidized form with the active site should be weaker than its non-oxidized form; and third, the non-oxidized form should stabilize strongly and close to the T1 copper atom. These findings align with theoretical frameworks proposed by other researchers(Xin et al., 2019), who have suggested that the migration of a single electron from the substrate to the enzyme involves a quantum tunnel from the most oxidizable atom of the substrate to the T1-copper atom in the enzyme. Consequently, the distance of this quantum tunnel plays a critical role in determining the effectiveness of the oxidation process. While interaction energy is often used as a predictor of enzymatic activity, our findings suggest that it represents only one aspect of the complex interplay between enzyme, substrate, and mediator. A more comprehensive understanding of these interactions, including the spatial positioning of substrates and the thermodynamic driving forces for product release, is necessary to fully optimize the degradation process and to design more efficient biocatalysts.

By combining experimental and computational approaches, our study demonstrates the importance of multidisciplinary research in addressing the challenges associated with antibiotic contamination. We hope these findings will inspire further studies aimed at developing more efficient and sustainable solutions for mitigating the risks associated with antibiotic resistance and for promoting a safer and more sustainable environment for future generations.

## Acknowledgements

1. J. M. Blamey and C. Toledo-Neira, thanks to DICYT Asociativo USACH 022242TN_DAS, by the Vicerrectoría de Investigación, Innovación y Creación of Universidad de Santiago de Chile. D. Muñoz-Gacitúa acknowledges support from a postdoctoral fellowship (POST-DOC_DICYT 022443BA) granted by the Vicerrectoría de Investigación, Innovación y Creación of Universidad de Santiago de Chile. The authors acknowledge and thank Fun-dación Biociencia for providing the computational resources that made this research possible.

## Credit authorship contribution statement

**Matias Donoso:** Experimental work, Writing – original draft. **D. Muñoz-Gacitúa**: Formal analysis, Conceptualization. **G. Salfate**: Analytical Methodology. **Carla Toledo-Neira**: Writing – review & editing and funding acquisition. **Jenny M. Blamey**: Conceptualization, Writing – review & editing, Validation, funding acquisition, and Supervision.

## AI assistance statement

During the preparation of this manuscript, the authors utilized a locally self-hosted large language model (LLM) to enhance grammar, sentence structure, and readability. To ensure data governance and maintain the confidentiality of unpublished research. All AI assistance was performed using self-hosted LLM infrastructure located at Fundacion Biociencia. All scientific content, methodology, data analysis, interpretations, and conclusions remain the original intellectual work of the authors, who take full responsibility for the accuracy and integrity of this publication.

## Data availability

The datasets generated and analyzed during the current study are available from the corresponding author upon request.

